# Hippocampal and Medial Prefrontal Cortical Maps Represent Episodes and Rules in a Common Task Space

**DOI:** 10.1101/2022.07.17.500349

**Authors:** Aditya Srinivasan, Justin S. Riceberg, Michael R. Goodman, Arvind Srinivasan, Kevin G. Guise, Matthew L. Shapiro

## Abstract

Memory helps us adapt to changing circumstances but needs guidance to retrieve relevant episodes. Episodic memory requires the hippocampus, the prefrontal cortex (PFC) guides memory retrieval, but how their representations interact is unclear. Using state-space analysis of neuronal spiking, we found CA1 and PFC activity within and between rats formed similar, low-dimensional, region-specific “shapes” representing different tasks tested in the same maze. Task shapes were organized by behaviorally salient variables including time and maze start and goal locations. PFC predicted CA1 representations when both regions were needed to solve a spatial memory task, but not in a cue approach task that required neither region. Task demands imposed common dimensions on CA1 and PFC maps whose topologies distinguished episodic and rule-related computations.

## Results

Episodic memory represents stimuli in spatial, temporal, and personal context^1,2^, and can guide choices by letting environmental features and internal goals evoke representations of available outcomes. When outcomes change, proactive interference^3^ impairs choices until evoked representations are updated. The neuronal mechanisms which update memory representations are unclear.

Coping with proactive interference requires the prefrontal cortex (PFC)^4,5^, and episodic memory requires the hippocampus^1,6^ in people and other animals. Tracking outcomes which change in space or time requires both the PFC and hippocampus^7–10^. For example, spatial reversal learning entails avoiding a previously rewarded location and instead approaching a previously unrewarded one. Intact rats readily learn initial spatial discriminations and reversals, inactivating the medial PFC (mPFC) impairs switching between goals^11^, and inactivating CA1 impairs both^12^.

mPFC and CA1 neurons respond to salient task variables. In reward locations, mPFC neurons generate firing patterns which track accumulating evidence about contingency rules^13^. mPFC neurons also discriminate spatial relationships between objects in an environment^7^, and different categories in visual classification tasks^8,9^. CA1 neurons discriminate external environmental variables including spatial location^14^, time^15,16^, auditory frequency^17^, odors, and taste^18–20^ along with internal variables such as social structures^21,22^. During reversal learning, which requires both structures, single mPFC and CA1 neurons respond to salient task variables including place, time, and goal. mPFC and CA1 ensembles each predict correct choices in single trials, and mPFC modulation of CA1 predicts learning speed^11^. While these observations suggest frontotemporal interactions update memory, they do not describe either the nature of mPFC or CA1 representations or their dynamics.

Neuronal representations can be rigorously described by analyzing the statistical structure of ensemble activity recorded from the behaving brain^23^. This approach describes sequential activity recorded in a neuronal population as successive points in a high-dimensional activity space^23–26^. Patterns of activity describe geometric shapes, analogous to the way a round-trip through three cities forms an approximate triangle on a road map. The geometry of activity spaces describes neuronal representations of motor control^27,28^, head direction^29^, and other task variables^23,30^, and provides a framework for bridging cognitive and neurophysiological levels of analysis.

Here, we used a state-space approach to analyze mPFC and CA1 ensembles from rats performing spatial memory and cue approach tasks in a plus maze^11^. We found both mPFC and CA1 activity spaces formed low-dimensional maps with dimensions corresponding to latent task variables including trial sequences, starting locations, and goal locations. Different neuronal subpopulations within brain areas formed similar maps even when recorded from different rats. mPFC and CA1 maps differed topologically, as did some maps of the spatial memory and cue approach tasks. mPFC maps consistently predicted CA1 map dynamics only when animals were learning to switch between spatial goals. The results suggest CA1 and mPFC contribute to flexible cognition by representing salient task features in maps with common dimensions and different topologies. CA1 maps separate representations of goal-directed spatiotemporal episodes, while mPFC maps combine episodes with shared goals.

### High-Dimensional Activity Spaces are Described by a Few Latent Dimensions

State-space analysis of neural activity produces high-dimensional matrices which estimate single unit and pairwise unit co-activity for each theta cycle in every trial of a recording session. A typical CA1 recording session of 60 units, 50 trials, and 16 theta cycles/trial is represented by an 1830-dimensional activity space. Each point indicates the activity state of the ensemble in one theta cycle. In this example, a trial would consist of 16 sequential points. The full set of points consists of the points from each trial. In the activity space, this set describes neuronal activity as a geometric and topological “shape” which represents brain activity associated with learning and memory performance. If the activity of single neurons in ensembles were independent, then describing representations accurately might require all the dimensions.

Because single neuron activity is constrained, e.g., by synaptic connectivity between pairs of neurons^31^, ensembles include redundant signals. Therefore, representations can be described accurately by fewer dimensions which maintain sufficient statistical variance. Using a non-linear dimensionality reduction method^30^, we found 4-6 latent dimensions accounted for 95% of the variance in CA1^23^ and mPFC activity spaces. The following sections refer to the low-dimensional representations as “embedded maps,” and the high-dimensional state spaces as “full maps” or activity spaces. Patterns suggested by embedded maps were quantified in full maps.

### Latent Dimensions of Activity Spaces Systematically Represent Task Variables

If the latent dimensions of the activity space correspond with task structure, then locations within the space should represent collections of task features, and paths through the space should correspond to behavior sequences. Three-dimensional embeddings of the high-dimensional activity spaces using MDS **(Fig. 1A-C, 2)** and t-SNE **(Fig. 1D)** showed neuronal activity organized along dimensions corresponding to journeys, time, and maze location **(Figs. 1, 2)**.

**Figure 1.**
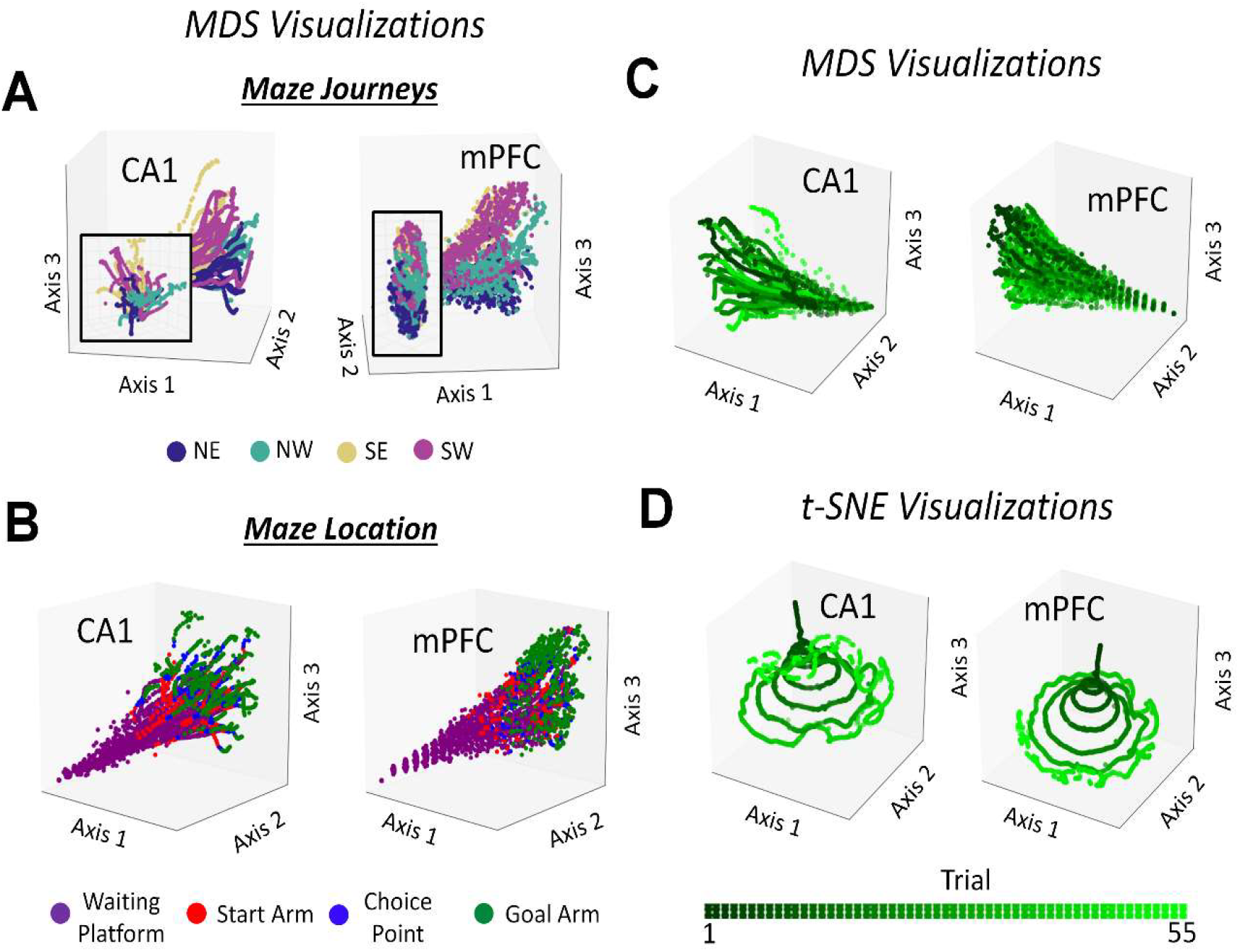
Maps Reveal Task Structure. Representative 3D images of low-dimensional embeddings of maps are shown via multidimensional scaling (MDS) and t-stochastic neighbor embedding (t-SNE). Each dot corresponds to the embedded neural activity during a single theta cycle. **(A)** MDS revealed journeys are segregated into different portions of the map in both CA1 and mPFC. Insets show the segregation of journeys at the base of the conical structure. **(B)** Alternate labeling of the MDS visualizations revealed neural activity in the waiting platform, start arm, choice point, and goal arm formed sequential, segregated clusters. Single trajectories through the map were temporally organized: neural activity in the waiting platform preceded start arm neural activity which preceded choice point and goal arm neural activity. **(C)** MDS embeddings do not show the sequential progression of trials through a testing session while **(D)** t-SNE embeddings do. Trials from the start of the session are shown at the apex of cone (black); those from the end of the session surrounded the base of the cone (light green).

**Figure 2.**
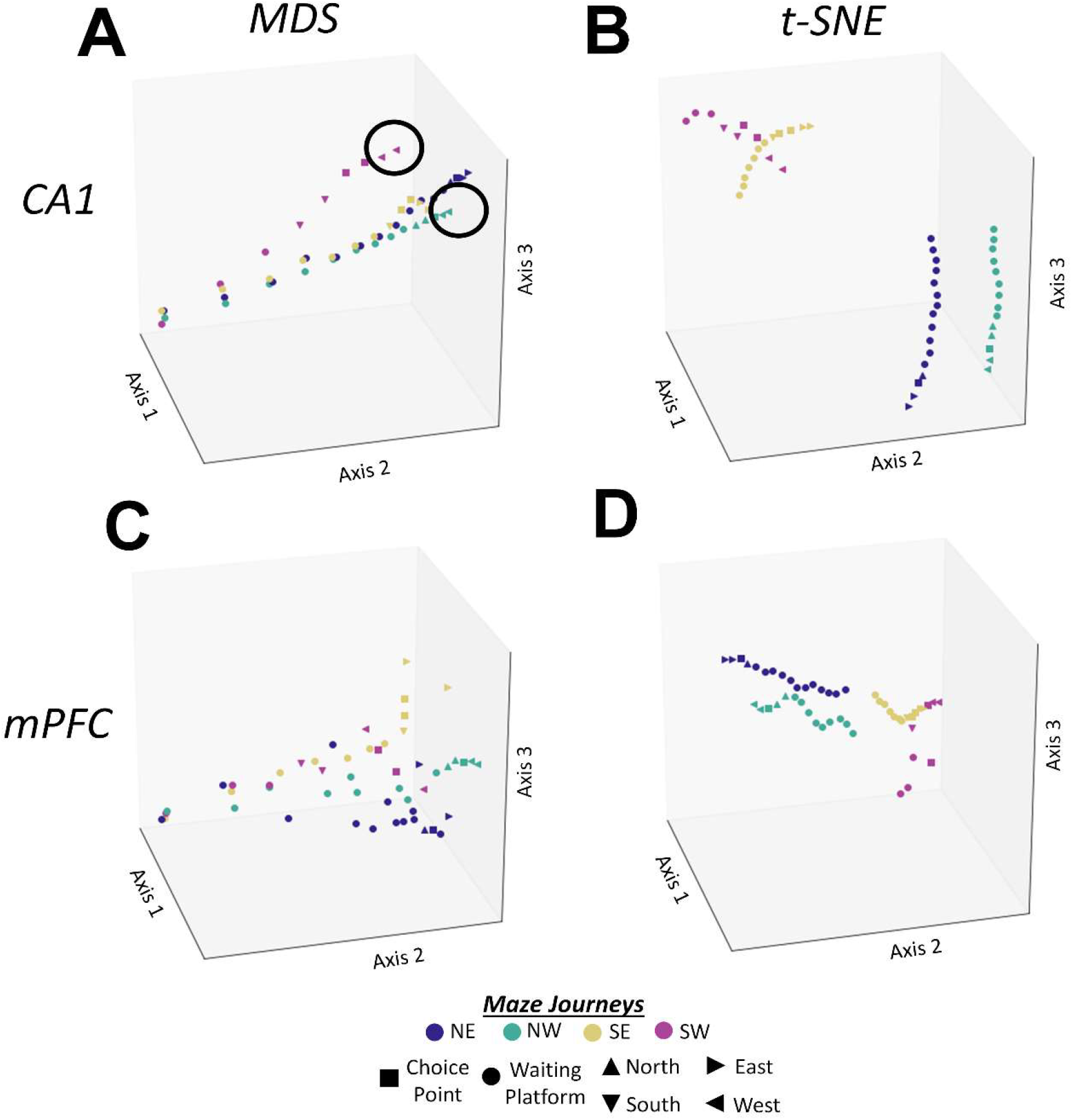
Single Trials Emphasize Differences Among Journeys and Embeddings. Differences between the features each embedding reveals are visualized by showing **(A-C)** CA1 and **(B-D)** mPFC neural activity from one “Northwest,” “Northeast,” “Southwest,” and “Southeast” journey through the plus maze. Coding locations were made more visible in single trajectories by showing every fourth point in each trajectory. **(A)** Circles indicate CA1 trajectories encode the same maze location, the “West” arm, in different activity space locations based on different journeys, i.e., the same goal reached from different starting locations. **(A-B)** MDS segregated the differences between starting and goal locations of single journeys; **(C-D)** t-SNE did not. Axis orientation was chosen to best visualize the trajectories.

Figure 1 shows examples of 3D visualizations of CA1 and mPFC maps of a full testing session; each dot shows the embedded location of neural activity in one theta cycle. Trajectories, filament-like sequences of dots, represent single trials as rats move from the waiting platform through the start arm, choice point, and goal arm. In Figure 1A, dot color indicates the types of *journeys* taken by the rat from start to goal, e.g., a trajectory of gold dots stretching from the lower left to the top right of the embedded CA1 and mPFC activity spaces corresponds to a SE journey **(Fig. 1A)**. The same trajectories are colored differently to indicate trial phase (waiting platform, start arm, choice point, goal arm) and trial number **(Fig. 1B, 1D** respectively). The overall shape of MDS and t-SNE embeddings of CA1 and mPFC activity spaces are roughly conical and exhibit spatiotemporal organization. MDS embeddings depict all trajectories originating from one corner of the map, corresponding to the waiting platform; trajectories diverge from there **(Fig. 1B)**. t-SNE embeddings further illustrate the sequence of trials along an axis orthogonal to an axis representing time within individual trials **(Fig. 1D)**. Examples of single trials in MDS and t-SNE embeddings better illustrate how trajectories through the activity space correspond with maze journeys **(Fig. 2)**. From the start of each trial, trajectories in the activity space diverged, even while the rat was on the waiting platform (circular dots, **Fig. 2A)**. By the end of corresponding journeys, identical maze positions occupied separate locations in the activity space (e.g., turquoise and magenta “West” triangles circled in **Fig. 2A)**.

### Maps are Conserved Across Neuronal Ensembles and Rodents within CA1 and mPFC

Embedded maps indicate CA1 and mPFC represent salient task features **(Figs. 1-2)**, but do not describe the variability of maps within or across rodents. At one extreme, maps could be determined purely by experience, and differ across ensembles recorded from the same brain region and animal. At the other extreme, maps constrained by species-typical networks could be similar across subpopulations within each brain region. To distinguish these possibilities, we measured the bottleneck distance between full maps formed by different ensembles recorded in the same task and brain structure. Full maps recorded in the same task and brain structure were more similar than expected by chance even when ensembles were recorded from different rats (Bottleneck Distance Mean ± SEM; *spatial memory task*: CA1 v. CA1 = 0.0020 ± 2.0*10^−4^, mPFC v. mPFC = 0.0022 ± 2.0*10^−4^; *cue approach*: CA1 v. CA1 = 0.0022 ± 7.3*10^−5^, mPFC v. mPFC = 0.0022 ± 7.3*10^−5^; **Fig. 3A-B)**. Rather than depending purely on experience or individual differences, ensemble representations are constrained by species-typical neuronal networks.

**Figure 3.**
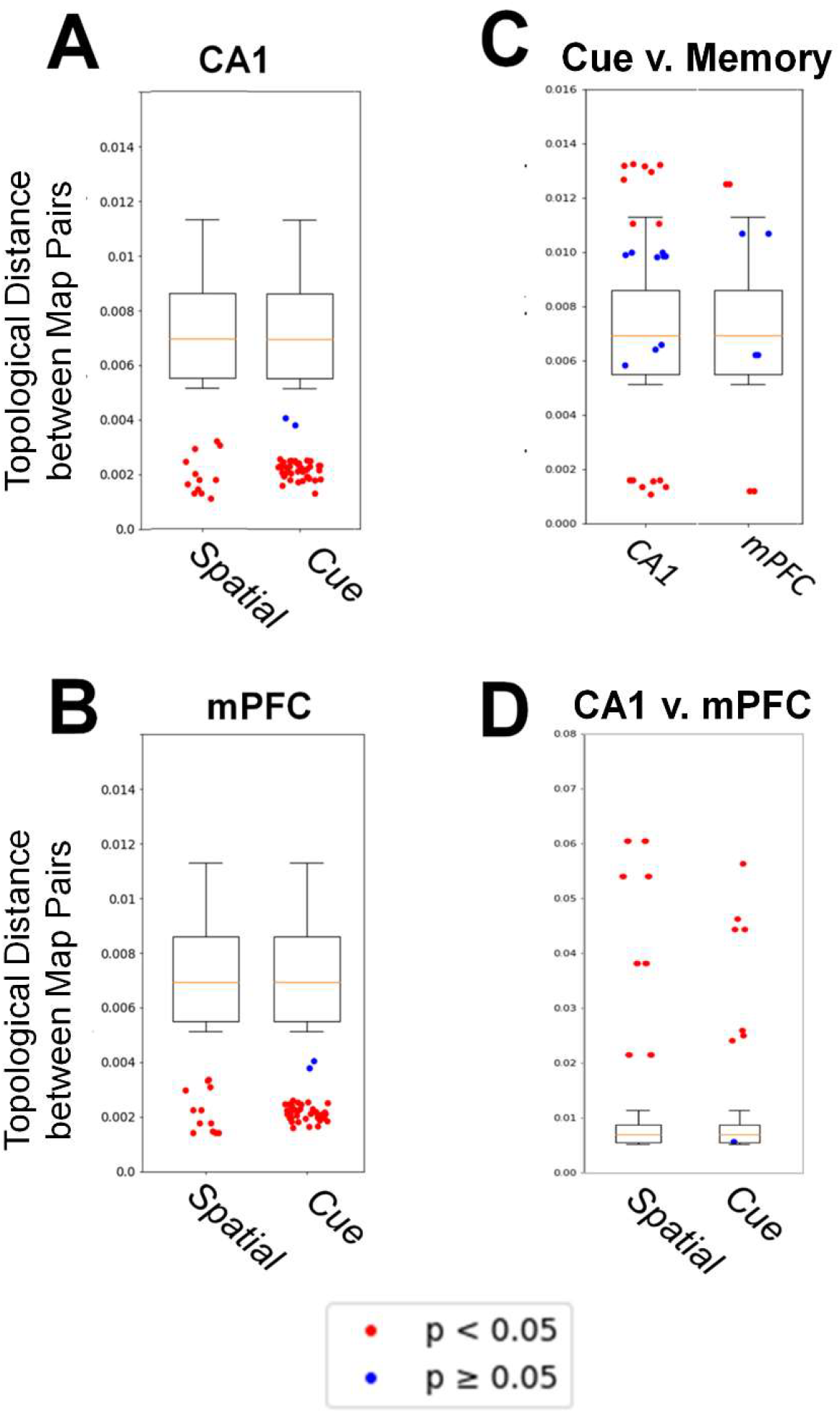
Similar Task-Specific Maps are formed by Different Groups of Neurons in CA1 and mPFC, Maps may be Shared Among Tasks, and Different Regions Construct Unique Maps. Maps constructed from neural activity from different populations of neurons recorded from **(A)** CA1 and **(B)** mPFC formed similar maps of the same task. **(C)** Maps constructed from only CA1 units and only mPFC units sometimes form maps which are more topologically similar than chance (red, below boxplot), as similar as chance (blue), or more dissimilar than chance (red, above boxplot). **(D)** Maps constructed from simultaneously recorded CA1 and mPFC units formed distinct maps of the spatial memory and cue approach tasks (*spatial memory*: 8/8 ensembles p < 0.05; *cue approach*: 7/8 ensembles, p < 0.05; permutation testing in all cases). **A-D** Dots represent individual topological distances between pairs of maps constructed from recordings of different populations (red, p < 0.05; blue, p > 0.05; permutation testing). The distribution of topological distances from randomly generated data is shown as a boxplot.

### CA1 and mPFC Maps of Identical Behavioral and Environmental Conditions Distinguish Task Demands

To determine how cognitive, behavioral, and environmental features modulated activity, we compared the same ensembles recorded in the spatial memory and cue approach tasks. Both tasks entailed identical overt responses to the same stimuli and differed only in reward contingency and cognitive demand. The spatial memory task required learning and remembering goal locations independent of a light cue; the cue task required rats to approach the light cue independent of location. If ensembles represent only environmental features and overt behavior, then representations of the two tasks should be similar. Alternately, if ensembles represent only cognitive demands, then the representations should differ markedly. The bottleneck distance between full maps showed some ensembles differentiated the tasks more than expected by chance (8 CA1, 2 mPFC ensembles), some represented the tasks as more similar than chance (8 CA1, 2 mPFC ensembles), and others as no different than chance (8 CA1, 4 mPFC ensembles; **Fig. 3C)**. As might be expected, CA1 and mPFC ensembles represented both cognitive and behavioral task features.

### CA1 and mPFC Ensembles Map Task Features with Different Topologies

Both mPFC and CA1 embedded maps represented salient task structure including goal-directed journeys, time, and place. If mPFC and CA1 circuits perform different computations^32^, however, then their full maps should differ. We therefore compared CA1 and mPFC full maps from pairs of simultaneously recorded ensembles using bottleneck distance. Simultaneously recorded CA1 and mPFC ensembles formed different full maps of the same task (permutation testing; Bottleneck Distance Mean ± SEM *spatial memory task:* 0.043 ± 0.0038, 8/8 ensembles, p < 0.05; *cue approach task:* 0.032 ± 0.0023, 7/8 ensembles, p < 0.05; **Fig. 3D)**. The different topologies suggest that though CA1 and mPFC both represent the same set of salient task features, each region computes different combinations of the task features’ variance.

### Different Circuits Parse Task Space: CA1 Maps Journeys, mPFC Maps Goals

Embedded maps show similarities and differences between CA1 and mPFC representations. Both structures represent spatiotemporal sequences of single trials as trajectories through maps. To quantify sequences within single trials, we compared the Mahalanobis distances between the centroids of each trial phase (waiting platform, start arm, choice point, and goal arm) for every trial in full maps. The distances between the centroids corresponded with journey sequences within trials, e.g., waiting platform centroids were closer to the start arm than to goal arm centroids. Trial phase sequences computed from full maps were indistinguishable from the actual behavioral sequence in individual trials across tasks and brain regions (PERMANOVA Testing; *Spatial Memory*: CA1: *R*_*3, 442*_ = 1, p = 0.01, mPFC: *R*_*3, 442*_ = 1, p = 0.01; *Cue Approach*: CA1: *R*_*3, 367*_ = 1, p = 0.01, mPFC: *R*_*3, 367*_ = 1, p = 0.01; **Tables 1, 2**).

**Table 1.**
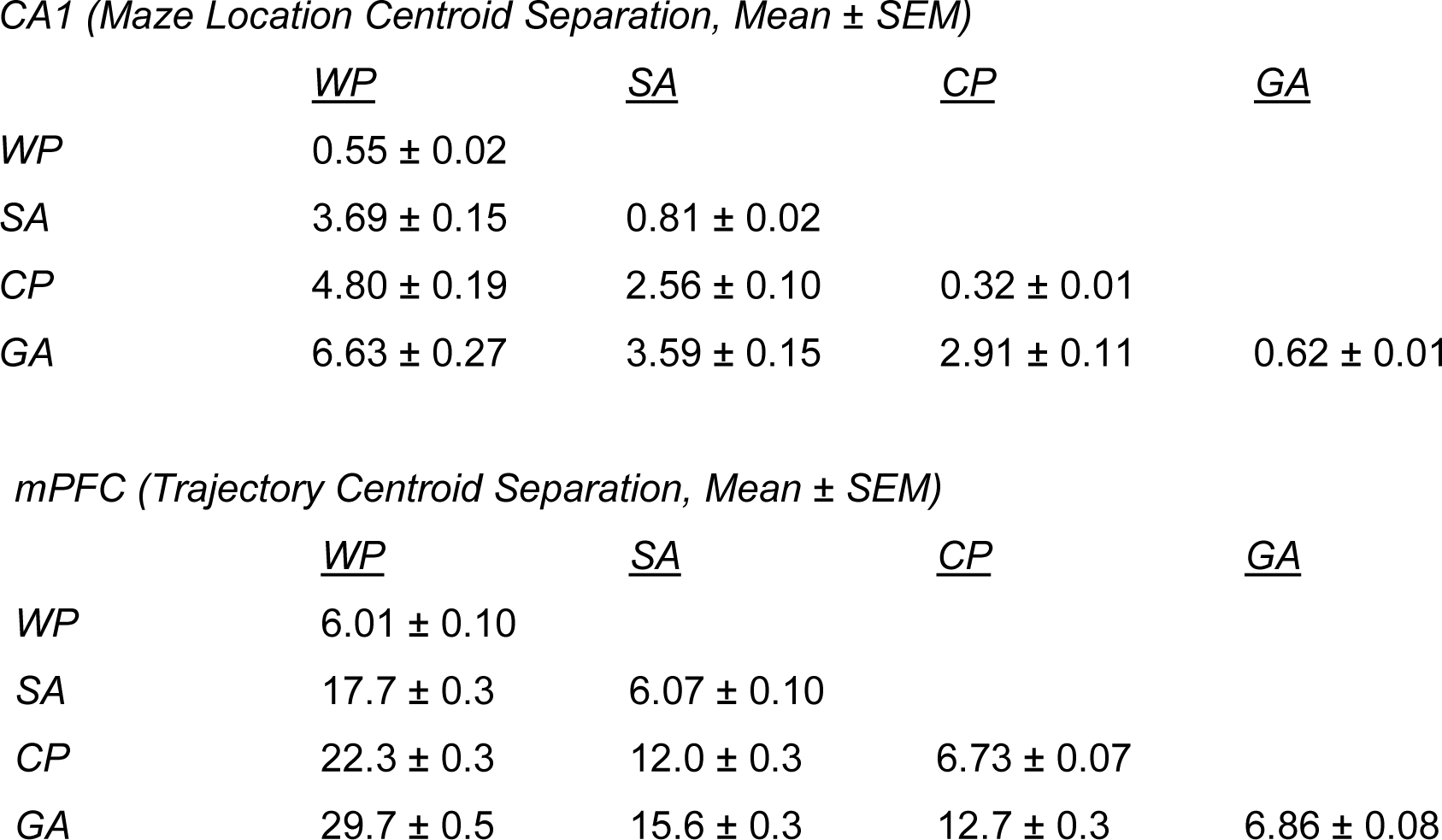
CA1 and mPFC Track the Sequence of Trial Epochs in the Spatial Memory Task. In the spatial memory task, CA1 and mPFC trajectories proceed through maze locations in sequential order from the waiting platform through the start arm, choice point, and goal arm. The location of each task phase is closest to itself across trials, e.g., the waiting platform is closest to the waiting platform (PERMANOVA of trial epoch; CA1: R_3,442_ = 1, p < 0.05, mPFC: R_3,442_ = 1, p < 0.05). WP – waiting platform, SA – start arm, CP – choice point, GA – goal arm.

|  | <u>WP</u> | <u>SA</u> | <u>CP</u> | <u>GA</u> |
| --- | --- | --- | --- | --- |
| WP | 0.55 ± 0.02 | | | |
| SA | 3.69 ± 0.15 | 0.81 ± 0.02 | | |
| CP | 4.80 ± 0.19 | 2.56 ± 0.10 | 0.32 ± 0.01 | |
| GA | 6.63 ± 0.27 | 3.59 ± 0.15 | 2.91 ± 0.11 | 0.62 ± 0.01 |

*mPFC (Trajectory Centroid Separation, Mean ± SEM)*
|  | <u>WP</u> | <u>SA</u> | <u>CP</u> | <u>GA</u> |
| --- | --- | --- | --- | --- |
| WP | 6.01 ± 0.10 | | | |
| SA | 17.7 ± 0.3 | 6.07 ± 0.10 | | |
| CP | 22.3 ± 0.3 | 12.0 ± 0.3 | 6.73 ± 0.07 | |
| GA | 29.7 ± 0.5 | 15.6 ± 0.3 | 12.7 ± 0.3 | 6.86 ± 0.08 |

**Table 2.** CA1 and mPFC Track the Sequence of Trial Epochs in the Cue Approach Task. In the cue approach task, CA1 and mPFC trajectories proceed through maze locations in sequential order from the waiting platform through the start arm, choice point, and goal arm. The location of each task phase is closest to itself across trials, e.g., the waiting platform is closest to the waiting platform (PERMANOVA of trial epoch; CA1: *R*_*3,442*_ = 1, p < 0.05, mPFC: *R*_*3,442*_ = 1, p < 0.05). WP – waiting platform, SA – start arm, CP – choice point, GA – goal arm.

|  | <u>WP</u> | <u>SA</u> | <u>CP</u> | <u>GA</u> |
| --- | --- | --- | --- | --- |
| WP | 0.83 ± 0.07 | | | |
| SA | 1.18 ± 0.05 | 0.86 ± 0.07 | | |
| CP | 2.57 ± 0.04 | 1.39 ± 0.06 | 0.82 ± 0.07 | |
| GA | 4.75 ± 0.06 | 3.58 ± 0.13 | 2.17 ± 0.09 | 1.21 ± 0.13 |

*mPFC (Trajectory Centroid Separation, Mean ± SEM)*
|  | <u>WP</u> | <u>SA</u> | <u>CP</u> | <u>GA</u> |
| --- | --- | --- | --- | --- |
| WP | 4.94 ± 0.23 | | | |
| SA | 8.95 ± 0.13 | 3.55 ± 0.18 | | |
| CP | 12.9 ± 0.2 | 7.75 ± 0.21 | 4.13 ± 0.41 | |
| GA | 29.2 ± 0.3 | 23.3 ± 0.4 | 9.75 ± 0.43 | 4.33 ± 0.34 |

t-SNE visualizations suggested both structures represent the temporal sequence of trials throughout testing sessions: trials at the start of the session were near the vertex of the cone; trials at the end of the session were near the base of the cone **(Fig. 1C)**. To quantify representations of trial sequences in each testing session, we calculated the centroid of each trial in the full map, calculated the distances between all pairs of trial centroids, and determined the shortest path through the distance matrix which included each centroid once and only once^33–35^. The reconstructed trial sequence matched the actual trial order better than chance in all ensembles for both tasks and brain regions (permutation tests, p < 0.05, 8/8 CA1 and mPFC ensembles, *Spatial Memory*: 446 trials, 55.8 ± 0.86 trials/session, reconstructed versus shuffled sequence errors, Mean ± SEM, *CA1*: 2.9 ± 0.28 vs. 25.6 ± 1.6; *mPFC*: 1.5 ± 0.47 vs. 25.5 ± 1.66; *Cue Approach*: 371 trials, 53.0 ± 2.82 trials per session, *CA1*: 1.75 ± 0.42, vs. 15.4 ± 2.55; *mPFC*: 1.50 ± 0.50 vs. 17.0 ± 0.28). CA1 and mPFC maps each represented temporal sequences of behavior within single trials and across all trials from the start to the end of the testing session.

MDS embeddings suggest CA1 better segregated journeys than mPFC **(Fig. 1A, 2**; e.g., compare the overlap between NW and SW journeys). In full CA1 maps, centroids of homogeneous journeys clustered closely together, and centroids of heterogeneous journeys were separated (ANOVA journey types; *Spatial Memory*: F_3,4904_ = 34.7, p < 0.01; *Cue Approach*: F_3,3062_ = 55.7, p < 0.01; **Fig. 4**). The distance between pairs of trajectories with a common arm was not statistically different from the distance between pairs of disjoint trajectories (e.g., NE, NW v. NE, SW; p > 0.05, Tukey HSD). In full mPFC maps, homogeneous journeys also clustered together, and different journey types were segregated. In contrast to CA1, journeys which shared the same goal arm were clustered more closely together than journeys with a shared start arm or disjoint journeys (ANOVA journey types; *Spatial Navigation*: F_3,4496_ = 35.8, p < 0.01; Tukey HSD, p < 0.05; *Cue Approach*: F_3,3062_ = 35.2, p < 0.01; Tukey HSD, p < 0.05; **Fig. 4**). In other words, mPFC maps generalized the starts, and differentiated the goals of journeys, whereas CA1 maps differentiated both starts and goals of journeys. These differences between CA1 and mPFC maps were consistent in both tasks.

**Figure 4.**
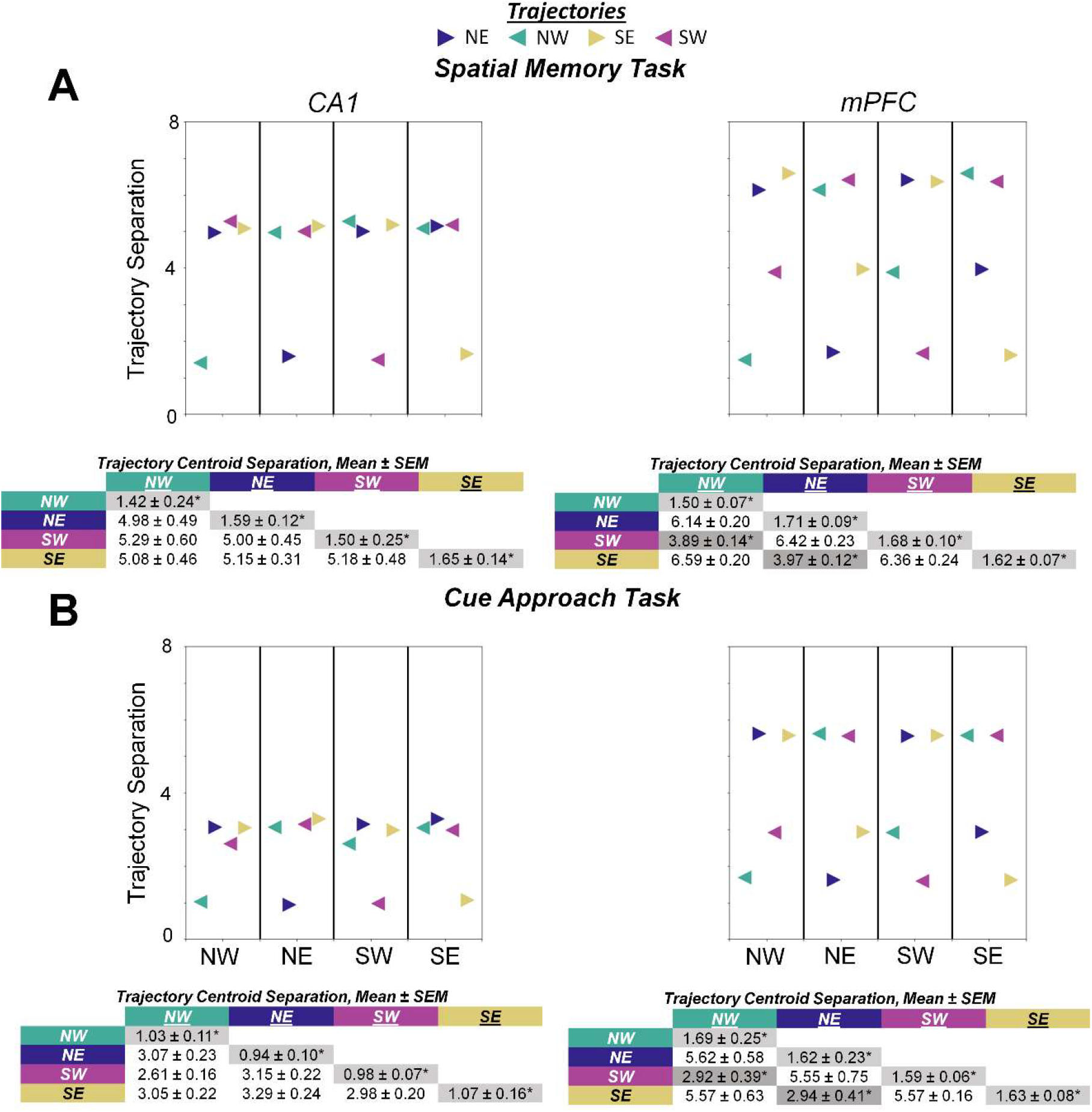
CA1 Maps Separate Trajectories by Starts and Goals, mPFC Maps Separate Trajectories by Starts and Group Trajectories by Goals. In both the **(A)** spatial memory and **(B)** cue approach tasks, CA1 strongly separated journeys based on starting and goal locations (ANOVA journey types; *Spatial Memory*: F_3,4904_ = 34.7, p < 0.01; *Cue Approach*: F_3,3062_ = 55.7, p < 0.01). In contrast, mPFC only separated journeys by starting locations and grouped journeys by shared goal locations (ANOVA journey types; *Spatial Memory*: F_3,4496_ = 35.8, p < 0.01; Cue *Approach*: F_3,3062_ = 35.2, p < 0.01). The colored symbols in the plots show the mean distance between journey types, SEMs are reported in the table. The shaded entries in the table show groups of journeys that differ significantly from other groups. Groups: 1) identical journeys: NW-NW, NE-NE, SW-SW, SE-SE, 2) homogeneous start arm: NW-NE, SW-SE, 3) homogeneous goal arm: NW-SW, NE-SE, 4) disjoint journeys: NW-SE, NE-SW, **\*** = Tukey HSD, p < 0.05 from other groups.

*mPFC Activity Modulates CA1 in the Spatial Memory, not the Cue Approach Task* The results so far show CA1 and mPFC maps are organized along similar dimensions corresponding to task features but have different topologies reflecting different coding hierarchies. CA1 and mPFC interact as rats perform spatial memory tasks^11,36^. To further investigate these interactions, we compared the temporal relationships between full CA1 and mPFC maps of each task using multivariate Granger prediction analysis. Multivariate Granger prediction analyses measures the extent to which changes in one map predict changes in the other map beyond what changes in each map predicts about itself. Each time series consisted of the sequence of centroids of journeys in the full maps for one ensemble from either CA1 or mPFC. In the spatial memory task, mPFC maps consistently predicted changes in CA1 maps (8/8 ensembles, permutation testing, p < 0.05), but CA1 maps rarely predicted changes in mPFC maps (1/8 ensembles, p < 0.05, permutation testing; Additional Variance Explained Mean ± SEM: mPFC → CA1 = 6.83 ± 0.716%, CA1 → mPFC = 2.73 ± 0.400%; **Fig. 5A)**.

**Figure 5.**
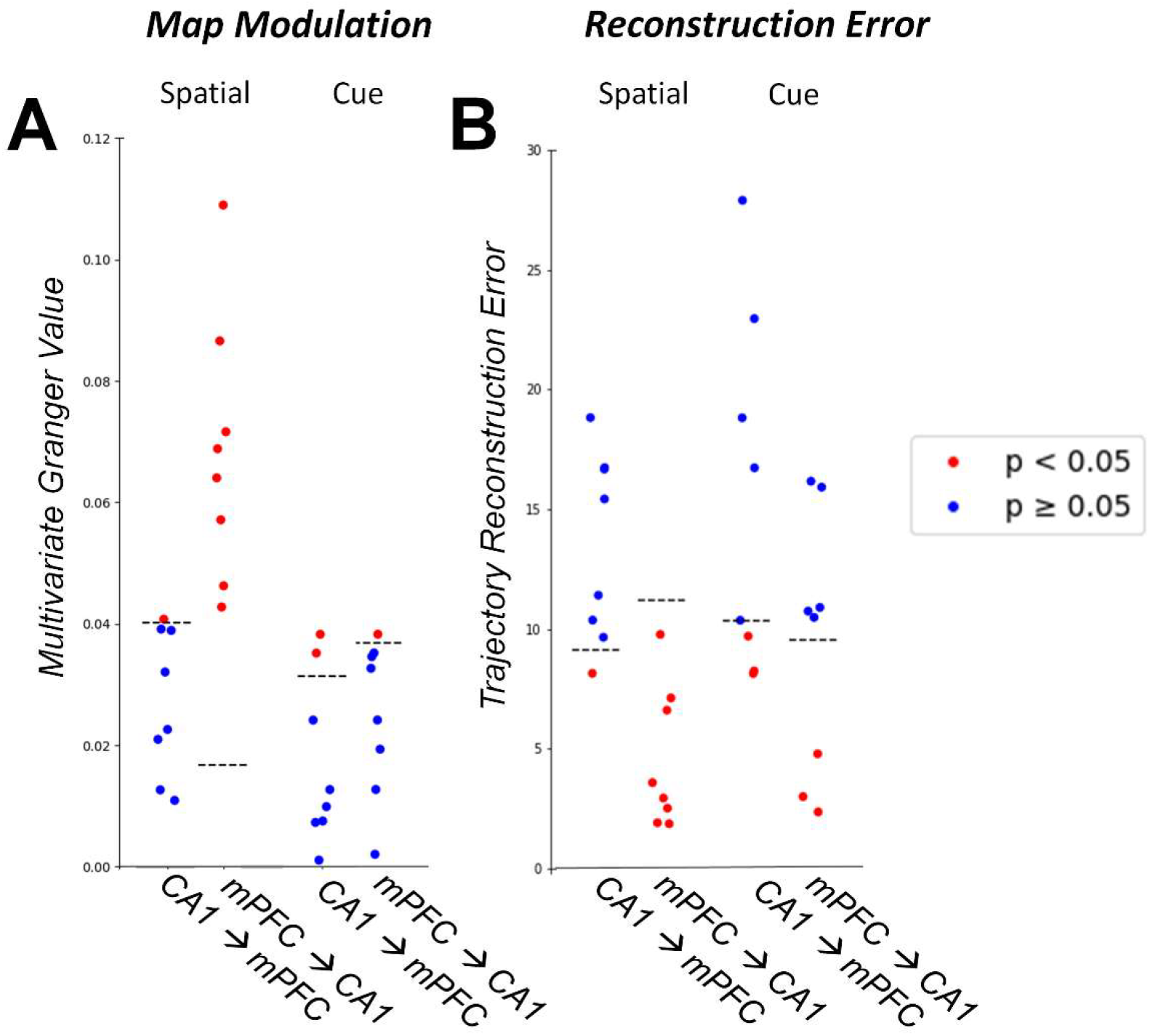
mPFC Maps Modulate CA1 Maps. **(A)** *Inter-Region Modulation:* During the spatial memory task, mPFC maps consistently modulated CA1 maps, while CA1 ensembles rarely modulated mPFC maps (CA1 → mPFC = 1/8 ensembles, p < 0.05, permutation testing, mPFC → CA1 = 8/8 ensembles, p < 0.05, permutation testing). During the cue approach task, maps rarely modulate one another (CA1 → mPFC = 2/8 ensembles, p < 0.05, permutation testing, mPFC → CA1 = 1/8 ensembles, p < 0.05, permutation testing). Frontotemporal interactions differed between tasks (*spatial memory v. cue approach*: Fisher Exact Test = 0.026, p < 0.05). Each dot shows the Granger value for one testing session. Black dashed lines represent the upper bound of the 95% confidence interval of the shuffled Granger values. **(B)** *Reconstructing Map Trajectories:* In the spatial memory task, the location of single trials in mPFC maps consistently predicted the location of single trials in CA1 maps, while CA1 rarely predicted the location of single trials in mPFC maps (CA1 → mPFC = 1/8 ensembles, p < 0.05, permutation testing, mPFC → CA1 = 8/8 ensembles, p < 0.05, permutation testing). In the cue approach task, neither structure formed maps that consistently predicted the location of single trials in the other structure’s map (CA1 → mPFC = 3/8 ensembles, p < 0.05, permutation testing, mPFC → CA1 = 3/8 ensembles, p < 0.05, permutation testing). Frontotemporal map interactions significantly differed between the tasks (*spatial memory v. cue approach*: Fisher Exact Test = 0.57, p > 0.05). Individual dots represent average error in trial location estimation, i.e., reconstruction error, for a session. Black dashed lines show the lower bound of the 95% confidence interval of the shuffled reconstruction errors. Red, p < 0.05; blue, p > 0.05, permutation testing.

Neither CA1 nor mPFC maps consistently predicted changes in the other in the cue approach task (permutation testing; Additional Variance Explained Mean ± SEM: mPFC → CA1 = 1.66 ± 0.340%, 1/8 ensembles, p < 0.05; CA1 → mPFC = 3.98 ± 0.592%, 2/8 ensembles, p < 0.05; **Fig. 5A**). The map dynamics are consistent with prior results comparing sequential changes in CA1 and mPFC ensembles^11^, verifying that changes in mPFC maps predict changes in CA1 maps, not vice versa. The map dynamics further show frontotemporal interactions occur during the spatial memory task which requires both structures, and not the cue approach task which requires neither. Together, the results suggest mPFC dynamics update memory representations by modulating paths through CA1 memory maps.

The multivariate Granger statistics analyzed mPFC-CA1 interactions over time across trials. Because the maps described here represent activity during every theta cycle, we reasoned they could reveal “instantaneous” frontotemporal interactions which modulate map structure. In other words, the location of a point in a mPFC full map should predict the location of a corresponding point in a simultaneously recorded CA1 full map. We used canonical cross correlation to predict the location of the centroids of individual trials in corresponding mPFC and CA1 maps. mPFC trial centroids consistently predicted the location of corresponding CA1 centroids during the spatial memory task (permutation testing; Reconstruction Error True v. Shuffled, Mean ± SEM: mPFC → CA1 = 4.53 ± 1.00 v. 24.5 ± 0.085, 8/8 ensembles, p < 0.05; **Fig. 5B**). In contrast, mPFC trial centroids inconsistently predicted the location of CA1 centroids in the cue approach task (permutation testing; Reconstruction Error True v. Shuffled, Mean ± SEM: mPFC → CA1 = 9.45 ± 1.85 v. 23.6 ± 0.090, 3/8 ensembles, p < 0.05; *spatial memory v. cue approach*: Fisher Exact Test = 0.026, p < 0.05; **Fig. 5B**). CA1 trial centroids did not reliably predict the location of corresponding mPFC centroids in either task (permutation testing; Reconstruction Error True v. Shuffled, Mean ± SEM; *Spatial Memory:* CA1 → mPFC = 13.7 ± 1.36 v. 16.9 ± 0.049, 1/8 ensembles, p < 0.05, *Cue Approach:* CA1 → mPFC = 15.7 ± 2.55 v. 24.7 ± 0.091, 3/8 ensembles, p < 0.05; *spatial memory v. cue approach*: Fisher Exact Test = 0.57, p < 0.05; **Fig. 5B**). CA1 representations are reliably predicted by mPFC activity only during the spatial memory task when both structures are required for learning and memory performance.

## Discussion

Neuronal population activity recorded simultaneously from mPFC and CA1 as rats performed spatial memory and cue approach tasks were described as >1000-dimensional activity spaces. Each point in the space defined the spiking of single units and co-activity of pairs of units in each theta cycle. Dimension reduction of the spaces showed salient task features were represented in approximately six-dimensional maps framed by common spatial, temporal, and behavior-related axes. Within maps, single trials formed smooth trajectories sequentially progressed through the maps as recording sessions proceeded and overlapped when rats performed similar goal-directed journeys. Similar maps were formed by different ensembles recorded from the same brain region and task. When task contingencies changed, some ensembles formed different maps. Map properties shown by the reduced maps were quantified in the full, high-dimensional activity spaces, and confirmed mPFC and CA1 representations differed. CA1 maps separated episodic journeys, while mPFC grouped journeys to the same goal and separated journeys to different goals. When rats performed a spatial memory task that required both structures, locations of mPFC activity predicted the location of simultaneously recorded CA1 activity in maps. When rats performed a cue approach task which entailed the same overt behaviors in the same environment but did not require either structure, map locations were unrelated. The common dimensions and different topologies of mPFC and CA1 maps suggest a general approach for analyzing the mechanisms that link neuronal computations across brain networks.

Previous studies assessed activity transition probabilities in CA1 ensembles and found geometric representations of task variables, e.g., maze location, evidence, and time^23^. Using a different method to define activity spaces^24,25^, the present results provide converging evidence that cognitive maps^37^ are computed by the hippocampus^2^, and these CA1 maps not only represent two-dimensional spatial location^38^ but also time, salient task demands, and space^23,37^. These additional variables help cognitive maps account for the full range of hippocampal contributions to episodic memory and cognition, including prospection^39^.

The sequences of neural activity observed in the embedded CA1 maps are consistent with the fast replay of firing sequences that accompany task performance^40–43^. Trajectories through the maps decoded time and behavioral goals as well as spatial locations. Therefore, fast replay sequences could be traversals along past trajectories within a multidimensional map. Further, hippocampal pre-play, sequences of neuronal activity observed both before and after rats first explore an unfamiliar environment^44^, could be traversals along potential trajectories in maps constrained by common inputs and synaptic connectivity.

The results also show hippocampal cognitive mapping is supported by prefrontal cortex. CA1 and mPFC both constructed maps framed by the same spatial, temporal, and behavioral dimensions. Like the hippocampus^2,11^, mPFC represents events as locations in a multidimensional space. Proximity in mPFC spaces, like in CA1 spaces, indicated similar relationships among salient task features, including the start and end of behavioral episodes.

CA1 distinguished identical maze locations in different journeys, signaling pending goal choices before, and past starting locations after rats traversed the choice point in the plus maze. Unlike CA1, mPFC maps separated journeys only by goal and generalized pairs of journeys which followed the same rule to the same goal^11,13^. Different ensembles of mPFC and CA1 neurons each formed similar maps, suggesting that circuit properties constrain topological representations of a given task. These circuit-specific representations may reflect *a priori* computations determined by different afferent inputs, intrinsic connectivity, and synaptic plasticity which support each region’s neuropsychological function. From this view, representations in different brain regions could form from either multiple map interactions or a distributed, brain-wide map which predicts the outcome of potential actions by combining multiple statistical computations.

Additionally, the maps represent both the overlap and disjunction of salient task features^23^. For example, the only difference between the cue approach and spatial memory task presented here was the reward contingency, signaled by a cue light or by remembering recently rewarded locations^11^. Partial remapping^45^, or other modulations of neural codes^44^ could support the solution of similar, but not identical, tasks^46^. When reward contingencies are altered while motivation state and the external environment remain constant, different subsets of ensembles likely represent one or another source of information, or combinations of the different sources. The distribution of ensembles that formed statistically distinct, similar, or indistinct maps in the cue approach and spatial memory tasks (**Fig. 3**) could reflect a pseudo-random sampling of neurons that tracked each information source. Further investigation is needed to determine if latent variables needed for solving one task can be adapted to another task, e.g., to promote learning sets or general schemas.

Recent computational work has investigated how representations of knowledge inform behavior in models of either reinforcement learning^47^ or predictive coding^11,48–50^ contexts. The salient task features extracted by these modeling techniques may concur with the latent variable structure identified in maps defined by state space analysis.

We found that the maps had a conical structure. Journeys diverged over time and were represented by trajectories which further separated as they approached the base of the cone. The non-conical structures formed by shuffled or randomly generated data suggest the conical structure is not a computational artifact^51^. Therefore, more work is required to better understand how activity maps correspond with the latent features discovered by modeling techniques, and how the structure of low-dimensional manifolds relates to memory representations and behavior.

Overall, the present results suggest that several brain regions represent learned knowledge geometrically, as multidimensional maps with features corresponding to relevant task features. The interplay between representations from multiple brain regions may construct brain-wide geometric representations of learned knowledge.

## Online Methods

Behavioral physiology experiments^11^ were performed in accordance with Institutional Animal Care and Use Committee guidelines and those established by the National Institutes of Health. Rats (n = 7) were trained on an elevated plus maze to perform a spatial memory task that requires both CA1 and mPFC function^11,52^. Each *testing session* was divided into four contingency *blocks* that required the rat to find chocolate sprinkles in either the “East” or “West” goal arm across repeated trials. In each trial, a rat was placed in either the “North” or “South” start arms, chosen pseudo-randomly, and learned to enter one goal to receive reward. Testing sessions began with an *initial discrimination*, during which animals learned which goal arm contained food by trial and error. After the animal entered the rewarded arm in 10/12 trials, the contingency block ended, and the other goal arm was rewarded in a *reversal*. During a single session of testing, rats completed an *initial discrimination* and 3 *reversals*. The rats were also trained in a cue approach task, wherein the animals found reward by approaching the goal arm indicated by a light cue. Neuronal ensembles were recorded as rats performed the spatial memory and cue approach tasks on the same day. Single unit activity from CA1 and mPFC were recorded simultaneously in three rats; in four other rats, only CA1 units were recorded^11^. The recorded ensembles of single unit activity were binned by theta cycles (∼125 milliseconds) and used in the following analyses. The results below are based on 8 simultaneously recorded CA1 and mPFC ensembles (CA1: mean = 61 units/ensemble, 488 units total; mPFC: mean = 28 units/ensembles, 224 units total), and 16 ensembles of only CA1 units (mean = 61 units/ensemble, 976 units total).

### Data and Code Availability

All code used here was developed in Python 3.7, Anaconda 2019.07 using the Jupyter IDE, MATLAB (R2019b, Mathworks), and several open-source packages^53–56^. Code and example data is available online (https://github.com/learningandmemorylab/Maps) and by request.

### Modeling Neural Activity using Manifolds

Multi-unit neuronal activity during a time interval can be described as a single point within a high-dimensional space. Each coordinate axis of this space corresponds to the activity of a single neuron. During behavior testing, ensemble activity can be described as a path, or trajectory, through the space.

If neuronal activity is constrained by task demands, then trajectories through the high-dimensional activity space may, in turn, be constrained by lower-dimensional sub-manifolds corresponding to salient task features. Discovering the structure of these geometric representations requires methods that accurately reconstruct these low-dimensional spaces from multi-unit activity. Ideally, these methods 1) discover nonlinear structure, 2) capture temporal relationships between states (successive points in the high-dimensional cloud of neural activity), 3) recover topological and geometric properties of the manifold, and 4) provide explicit mappings between neural activity and coordinates in the low-dimensional space^30^.

We used state space analysis of spike cross correlations (SSASC) which satisfied the above criteria.

### State Space Analysis of Spike Cross Correlations (SSASC)

We estimated unit activity and pairwise unit interactions using Shimazaki’s state space modeling^24,25^. Shimazaki’s state space model takes as input an R-tuple of spike times recorded from N neurons simultaneously^24^; hereafter, each repeated recording of the spike times is termed a trial. To analyze activity patterns of the recorded neurons, the parallel spike sequences are discretized into T time bins with bin size Δ, representing the population activity by a set of binary variables (either 1 or 0). For neurons n = 1, 2, …, N, time bins t = 1, 2, …, T, and trials r = 1, 2, …, R, the neural activity can be represented as the binary variable X_r,t,n_, where X_r,t,n_ = 1 if neuron n spiked during time interval t during trial r, else X_r,t,n_ = 0. Therefore, the entire recorded dataset can be described as a R × T × N dimensional binary array.

Dynamic population activity was modeled by a state-space approach which consisted of an observation model and a state model. The observation model specified the probability distribution of population activity patterns using state variables, while the state model described how those state variables change. The observation model for pairwise interactions (2^nd^ order) of neurons was defined as:

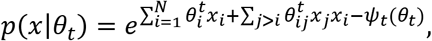

where x = [x_1_, x_2_,…x_N_] is a random vector of unit activity, *ψ*_*t*_(*θ*_*t*_) is a log normalization term, and θ_t_ refers to all the natural parameters of an exponential family distribution which are comprised of: 1) θ_i_ – the contribution of a single unit’s activity to the overall spike train and 2) θ_ij_ – the contribution of the interaction between units i and j to the overall spike train. Triplet, quadruplet, etc. interactions can be accounted for by increasing the expansion order terms in the above model (e.g., triplets: θ_ijk_).

The state model that considers the dynamics of a latent state θ_t_ is described by a random walk:

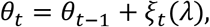

where *ξ*_*t*_ is a random vector drawn from a multivariate normal distribution *N*(0, Q). Q is a diagonal covariance matrix, and it is assumed the entries of the diagonal of the inverse matrix Q^-1^ are given by a scalar λ that determines the precision of noise for all elements within Q. The initial state θ1 is modeled as a normal random variable with mean μ and standard deviation Σ, i.e., *p*(*θ*_1_) = *N*(*µ, Σ*).

Given the data X (R × T × N above), the goal is to jointly estimate the posterior density of the latent states and optimal noise precision λ. Denoting hyperparameters of the model as ω = (λ, μ, Σ), the posterior density of the state process can be written as:

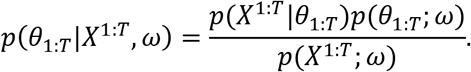

The first factor in the numerator is constructed from the observation model and the second factor from the state model. This posterior density can be constructed by approximating it by a Gaussian distribution, and depends on the choice of parameters w; the optimal w maximizes the marginal likelihood, i.e., evidence, that appears in the denominator given by:

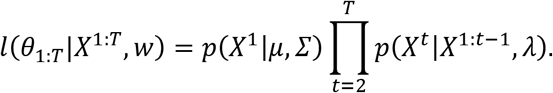

This optimization is achieved using an iterative approach that combines an expectation maximization algorithm with Bayesian recursive filtering and smoothing algorithms. In the resulting tridiagonal covariance matrix, the diagonal corresponds to the activity states of neurons 1 to N, and off-diagonals correspond to pairwise unit interactions, shown as:

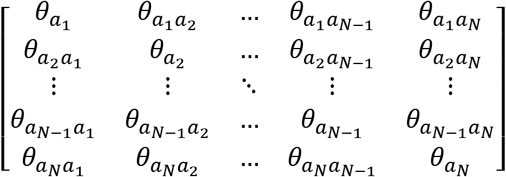

where 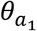 describes the activation state of unit 1, 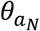 describes the activation state of unit N, and 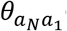 describes the pairwise coactivity of units 1 and N^25^.

The above matrices describe the connectivity state of all recorded neurons for every theta cycle while behavior is ongoing and are used in the subsequent analyses.

### Generating Artificial Neural Activity to Compare Between Maps

To interrogate the consistency of maps, we compared the bottleneck distances between the maps generated from real data and from artificial pseudorandom data. By comparing maps generated from the random data against maps generated from the real data, we ensure observed differences in maps of electrophysiological data are not due to variance in the modeling methods, e.g., differences in the fitting of the SSASC posterior density of latent states.

For each recorded ensemble, in each brain region (CA1 and mPFC), we first generated artificial spike trains by modeling each unit’s spike sequence as a Poisson process – each process has only one parameter which describes the mean firing rate of the unit over a behavioral epoch^57,58^. These processes generate spike trains which preserve the overall number of spikes per unit across all units, but do not preserve temporal interrelationships between unit spiking activity.

We then generated a second artificial dataset by randomly shifting 125 millisecond (∼1 theta cycle) population activity vectors in both CA1 and mPFC either forward or backwards in time^59^ preserving the inter-spike intervals within and between units, but not preserving trajectories through the neural activity space. This second method preserves the overall geometry of the space, unlike the first approach, but does not preserve the sequential organization of individual activity states within a trajectory.

We used both approaches to generate artificial ensembles from each recorded ensemble to compare the similarity of maps constructed from real recorded electrophysiological data as compared to the similarity of maps constructed from artificially generated data.

### Topological Data Analysis (TDA)

TDA is the branch of mathematics that applies tools from algebraic topology to extract information about the large-scale geometric structure of data. Real high-dimensional data is almost always sparse, and the relevant features often manifest in lower dimensions. TDA presumes the “shape” of data matters and attempts to provide a precise characterization of this “shape” via computation of coarse geometric descriptors.

To extract meaningful geometric information from recorded neural activity, we constructed a pairwise Euclidean distance matrix from the array of states, i.e., a map or activity space, estimated by SSASC. Either input distance matrix was then used to construct a Vietoris-Rips complex, a method of forming a topological space from distances in a set of points. Computing the Vietoris-Rips complex of a space M over a range of scales δ results in a so-called filtration of simplicial complexes. Computing simplicial homology for this filtration yields a multiset of “birth and death times” for homology classes within the growing simplicial complex. A graphical representation of the growing simplicial complex is given by a persistence diagram consisting of a multiset of points:

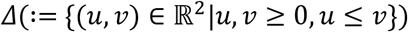

Persistence diagrams and barcodes describe the geometric and topological properties of high-dimensional data by enumerating the number and type of lower dimensional “holes” in the space. To measure topological differences between a pair of two high-dimensional activity spaces (point clouds), we generated a persistence diagram for each and computed the bottleneck distance between them^60^. The bottleneck distance measures the minimum changes required to transform one point cloud into another, i.e., by moving the fewest points the least distance. Formally, the bottleneck distance *W* between two points clouds *X* and *Y* is:

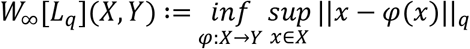

where q ≤ ∞ and the infimum is taken over all bijections X → Y. The bottleneck distance is typically calculated independently for each dimension d < ∞ ^61^. Typically, most differences lie in the 0, 1, and 2-dimensional homology spaces. As such, we can then define a combined bottleneck distance as:

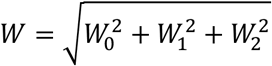

where 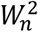 refers to the squared bottleneck distance between the n-dimensional persistence diagrams X and Y. We use this combined bottleneck distance W for further analyses, and compare W generated from real maps against W generated from pairs of maps generated from artificial data (described above) via permutation testing. In other words, bottleneck distances quantitate how much topologies differ.

### Estimating Intrinsic Dimensionality of Maps

The neural activity we describe as maps can be described equivalently as point clouds in high-dimensional coordinate systems. Each coordinate axis of the systems corresponds to the activity of a single recorded unit or co-activity between a pair of units. For an ensemble of 60 units, this results in an 1830-dimensional space. Such high-dimensional spaces are hard to understand intuitively. To estimate the variance in the high-dimensional spaces within a smaller number of dimensions, we used a non-linear dimensionality reduction technique which quantifies the additional variance explained by successive dimensions^30^. This approach estimates the minimum number of dimensions that optimally describe the high-dimensional point cloud defined by neural activity. For each dimensionality d, the dimensionality score G(d) was computed using the original data points X, their low-dimensional manifold coordinates Y, and their reconstructions 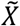. The score implements a tradeoff between two terms – the similarity between X and 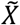, and the similarity of Y to a uniform d-dimensional ball. The dimensionality score is defined as *G*(*d*) = *−max* (*G*_1_, *G*_2_) where G_1_ is the 95^th^ percentile of the reconstruction errors 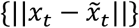 and G_2_ is the 95^th^ percentile of the sets of distances {H(±v_j_, Y)} across all v_j_, 1 ≤ j ≤ d where v_j_ is the standard basis vector in ℝ^d^ and:

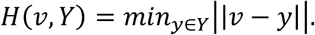

As d increases, G_1_ decreases while G_2_ increases. The optimal manifold dimensionality, d, maximizes G(d). Changing several parameters of the definition of G(d), including the percentiles or replacing percentiles by mean or median, gave equivalent estimates of optimal manifold dimensionality^30^.

### Mapping High-Dimensional Spaces into Low-Dimensional Ones

Lower dimensional representations (embeddings) help visualize the similarities and differences between neural activity states and trajectories, sequences of states which accompany behavior. After determining the optimal dimensionality of the data, we used two dimension reduction techniques that emphasize global and local distances among points: multidimensional scaling (MDS) and t-stochastic neighbor embedding (t-SNE). MDS, a deterministic approach, embeds high-dimensional points by maintaining the relative distance among all pairs of points^62^. MDS takes as input a matrix consisting of the pairwise distance, or dissimilarity, of points in the high-dimensional space and minimizes strain, a deterministic loss function, to map the points directly from a higher-dimensional space to a lower-dimensional one^62^. The output point cloud preserves the relative distance between all pairs of points and maintains local geometric and topologic properties across dimensionality^63^. t-SNE also takes as input a matrix consisting of the distances between pairs of high-dimensional points but instead uses the probability of neighborliness to assign points to their location in a lower-dimensional space. t-SNE embeddings minimize the difference between the probabilities that pairs of points are neighbors across dimensionality. Concretely, the difference between points in the high dimensional space defined, e.g., by Euclidean distance, is used to compute the probability of neighborliness given the empirical distribution of distances. Points are assigned to lower dimensional locations by minimizing the Kullback-Liebler cost function matching neighborliness probabilities in the low-dimensional, embedded data with the high-dimensional data^64^.

### Comparing CA1 and mPFC Encoding of Journeys

To quantify the different activity patterns in CA1 and mPFC indicated by MDS and t-SNE, we calculated the Mahalanobis distance^65^ between the centroids of trajectories corresponding to different journeys. To ensure the results were not affected by embedding techniques, we computed the centroids of trajectories (each *trajectory* corresponded to one *journey*) in the empirical, high-dimensional activity spaces. We used one-way ANOVA and Tukey’s HSD post-hoc testing to compare the distances between the centroids of homogenous journeys (NE-NE, NW-NW, SE-SE, SW-SW), journeys with common start arms (NE-NW, SE-SW), journeys with common goal arms (NW-SW, NE-SE), and disjoint journeys (NW-SE, NE-SW).

### Comparing CA1 and mPFC Encoding of Time

To quantify the encoding of behavioral sequences in CA1 and mPFC suggested by MDS and t-SNE, we calculated both the intra-trial and inter-trial Mahalanobis distances. As when quantifying CA1 and mPFC encoding of journeys, we computed the centroids of the trajectories in the empirical, high-dimensional activity spaces to avoid the influence of different embedding techniques. To calculate the intra-trial distance, we split each trajectory into four segments corresponding to time the animal spent in the waiting platform, start arm, choice point, and goal arm. We determined the centroid of each segment and calculated the Mahalanobis distance between all possible pairs of centroids (e.g., waiting platform to start arm). We used permutational multivariate analysis of variance testing, PERMANOVA^66^, to compare distances between the centroids of different maze locations, analogous to intra-trial time.

To calculate the inter-trial distance for each testing session, we computed the centroid of each trial’s trajectory and then found the pairwise Mahalanobis distance between all pairs of trajectories, resulting in a pairwise symmetric distance matrix. Using the Floyd-Warshall algorithm^33–35^, we found the shortest path through the distance matrix which included all trials in the testing session. We then compared the order in which trials were visited in this shortest path to the actual ordering of trials during the testing session – these were expected to match if the distance between the centroids of trials in the activity space is proportional to the order in which trials occurred on the day of testing, e.g., trials 1 and 2 are closer together than trials 1 and 10. We then shuffled the trial order labels in the testing session and repeated the above procedure 1000 times to generate a null distribution of trial ordering differences. We compared the trial ordering from the activity space and the null distribution via permutation testing to ensure the activity space’s trial ordering closely resembled the behavioral session’s trial order.

### Quantifying CA1-mPFC Map Temporal Interactions

To quantify sequential interactions between CA1 and mPFC maps, we first computed the centroid of each trajectory in the CA1 and mPFC maps in each testing session. The centroid calculation resulted in matrices of the form n x m where n is the number of trials during the testing session and m is the number of units and unit co-activity pairs. We then performed multivariate Granger prediction analysis^67^ on the multivariate time series consisting of trial centroids. Univariate Granger prediction analysis assesses the temporally directed relationship between two univariate time series by testing if the recent history of one time series predicts changes in another target time series beyond what the history of the target series predicts about itself. The Granger value quantifies this prediction as the log ratio of the residual variances for the model of only the target series and the model incorporating both the target and other time series. Higher Granger values indicate a greater amount of variance in the target series is explained by including the second time series^67,68^. Multivariate Granger assesses the temporally directed relationship between two multivariate time series by finding the elements of an input, multivariate time series which best predict changes in another target multivariate time series. This is accomplished by creating different subsets of the input multivariate time series, termed explanatory vectors, and finding which of these explanatory vectors best explains changes in the target time series. As in the univariate case, the Granger value quantifies this prediction as the log ratio of the residual variances for the model of only the target series and the model incorporating both the target and best-performing explanatory time series^67^.

To determine the statistical significance of Granger values, we compared each session’s Granger value to chance using permutation testing. After calculating the residual variance of the model which incorporated only the target time series, we shuffled the non-target time series. This shuffling eliminated the temporal correspondence between the two multivariate time series and tested the extent to which any increase in explained variance of the target was produced by the increased number of independent variables provided by including the non-target series^11^. Permutation tests compared the actual Granger values to the shuffled values, the null distribution.

### CA1-mPFC Interactions: Optimizing Model Selection

The dependent variable for each observation in the multivariate Granger prediction analysis was the SSASC parameters that described unit activity and co-activity during a trial. Independent variables were the unit activity and co-activity parameters in previous trials for the target time series or both time series. The number of the independent variables was determined by minimizing the Bayes’ Information Criterion (BIC) with respect to trial tenths. The BIC aids in model selection by balancing model fit while minimizing the number of parameters^11,69^. The BIC was calculated using:

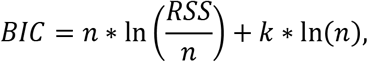

where n is the number of observations in the model, RSS is the residual sum of squares, and k is the number of parameters in the model which includes both time series.

### Quantifying CA1-mPFC Map Structural Interactions

To quantify structural interactions between CA1 and mPFC maps, we first computed the centroid of each trial’s trajectory in the CA1 and mPFC maps in each testing session. After the centroid calculation, each trial was described as a vector with length corresponding to the number of units and unit pairs recorded in the region during the testing session.

To determine if the structure of CA1 and mPFC maps could be used to predict the location of trajectories in the other structure, we used canonical-correlation analysis^70^ which determined a mapping from one structure’s map to the other’s. We determined these mappings from CA1 to mPFC and mPFC to CA1 for all but one trial in a testing session. We then used the mapping and held-out trial to predict the location of the trajectory corresponding to the held-out trial in the other map, leave-one-out cross-validation^71^. For example, if we held out trial 1, trial 1’s location in the CA1 map would be used to predict trial 1’s location in the mPFC map, and vice versa. We determined the reconstruction error, i.e., Euclidean distance, between the predicted location of the trajectory and the actual location of the trajectory. This procedure was repeated for all the trials in the testing session, and we averaged the reconstruction error across all trials in the testing session. To determine if the mappings were due to chance, we repeated the above procedure after shuffling the trial labels of the different trajectories 1000 times. This shuffling procedure generated a null distribution of reconstruction errors. For each session, we compared the average reconstruction error to the null distribution using permutation testing.

## Acknowledgements

This work was supported by the National Institutes of Health (NIMH 2R01MH073689, NIMH MH118297, and NIMH MH119523 to M.L.S.) and the NVIDIA corporation.

## Author Contributions

AS conceptualized the study, developed/validated the methodology and software used, and prepared the original draft. JSR and MRG helped develop the methodology and performed formal analysis. KGG acquired resources and curated the data. AS contributed to software development and validation. MLS helped conceptualize the study, provided supervision, acquired funding, and helped prepare the original draft. All authors contributed to draft review and editing.

